# Overabundant endocannabinoids in neurons are detrimental to cognitive function

**DOI:** 10.1101/2024.09.17.613513

**Authors:** Dexiao Zhu, Jian Zhang, Xiaokuang Ma, Mei Hu, Fei Gao, Jack B. Hashem, Jianlu Lyu, Jing Wei, Yuehua Cui, Shenfeng Qiu, Chu Chen

## Abstract

2-Arachidonoylglycerol (2-AG) is the most prevalent endocannabinoid involved in maintaining brain homeostasis. Previous studies have demonstrated that inactivating monoacylglycerol lipase (MAGL), the primary enzyme responsible for degrading 2-AG in the brain, alleviates neuropathology and prevents synaptic and cognitive decline in animal models of neurodegenerative diseases. However, we show that selectively inhibiting 2-AG metabolism in neurons impairs cognitive function in mice. This cognitive impairment appears to result from decreased expression of synaptic proteins and synapse numbers, impaired long-term synaptic plasticity and cortical circuit functional connectivity, and diminished neurogenesis. Interestingly, the synaptic and cognitive deficits induced by neuronal MAGL inactivation can be counterbalanced by inhibiting astrocytic 2-AG metabolism. Transcriptomic analyses reveal that inhibiting neuronal 2-AG degradation leads to widespread changes in expression of genes associated with synaptic function. These findings suggest that crosstalk in 2-AG signaling between astrocytes and neurons is crucial for maintaining synaptic and cognitive functions and that excessive 2-AG in neurons alone is detrimental to cognitive function.

## 1. Introduction

Endocannabinoids are the body’s naturally occurring bioactive lipid mediators that participate in a variety of physiological and pathological processes. Among them, 2-Arachidonoylglycerol (2-AG) is the most abundant endogenous cannabinoid and acts as a retrograde messenger that modulates synaptic transmission and plasticity at both inhibitory GABAergic and excitatory glutamatergic synapses in the brain via G-protein-coupled cannabinoid receptor 1 (CB1R) ^1–8^, which is also the target of Δ^9^-tetrahydrocannabinol (Δ9-THC), the main psychoactive ingredient in marijuana ^9^.

Accumulated evidence indicates that 2-AG signaling is crucial for maintaining brain homeostasis and protecting neurons in response to various stimuli or harmful insults ^10–13^, largely by resolving neuroinflammation ^12,14–18^. For instance, traumatic brain injury (TBI) triggers the release of 2-AG, and administering synthetic 2-AG significantly reduces TBI-induced edema and cell death while improving clinical recovery ^13,19,20^. These neuroprotective effects appear to stem from 2-AG-mediated mitigation of TBI-induced neuroinflammation ^17,20,21^. Similarly, 2-AG release is observed in the brain in response to the infusion of β-amyloid (Aβ) ^22^, and direct administration of 2-AG has been shown to protect neurons from proinflammatory and excitotoxic insults *in vitro* ^15,18,23^. These findings suggest that the release of endogenous 2-AG acts as a homeostatic defense mechanism, enhancing the brain’s resilience to injury, infection, or inflammatory stimuli, thereby maintaining brain homeostasis ^11,12,16^. Nevertheless, the released 2-AG is subject to rapid degradation by several enzymes, including monoacylglycerol lipase (MAGL), α/β hydrolase domain-containing protein 6 and 12 (ABHD6 and ABHD12), cyclooxygenase-2 (COX-2), and others ^12,24^. Among these, MAGL has been recognized as the primary enzyme responsible for degrading approximately 85% of 2-AG in the brain ^25–29^. The immediate metabolite of 2-AG is arachidonic acid (AA), a precursor of prostaglandins through catalytic activity of the enzymes COX-1/2 and of leukotrienes through the enzyme arachidonate 5-lipoxygenase (LOX). Certain prostaglandins (*e.g.*, PGE_2_) or leukotrienes (*e.g.*, LTB4) can act as proinflammatory mediators ^30–33^. This indicates that inactivating MAGL, which increases anti-inflammatory and neuroprotective 2-AG levels while reducing its proinflammatory and neurotoxic mediators, has ‘dual effects’ that help resolve neuroinflammation and provide neuroprotection against various harmful insults or pathological conditions ^12,15,18,23,34^. It is likely that 2-AG maintains brain homeostasis primarily by modulating synaptic transmission and plasticity as a retrograde messenger and by protecting neurons as an endogenous terminator of inflammation ^11,14,35,36^. Indeed, inhibition of 2-AG metabolism by pharmacological or genetic inactivation of MAGL has been shown to mitigate neuroinflammation, alleviate neuropathology, and improves synaptic and cognitive functions in animal models of brain disorders, including Alzheimer’s disease (AD) and TBI-induced AD-like conditions ^12,34,37–46^. Therefore, inhibiting 2-AG metabolism by inactivating MAGL has been proposed as a therapeutic approach for neurodegenerative diseases^12,36,37,47–51^.

However, the precise mechanisms underlying the neuroprotective effects of limiting 2-AG metabolism in neurodegenerative diseases remain to be fully elucidated. Notably, a recent study indicates that the neuroprotective effects of MAGL inactivation in mitigating TBI-induced neuropathology and preserving synaptic and cognitive functions are primarily due to the inhibition of 2-AG metabolism in astrocytes rather than neurons ^34^, suggesting a cell type-specific effect of 2-AG signaling in neuroprotection against brain trauma. Importantly, although 2-AG is widely recognized as a retrograde messenger in modulating synaptic transmission and plasticity ^1–8^, the role of 2-AG metabolism in long-term synaptic plasticity and cognitive function remains unclear. To better understand 2-AG metabolism in brain function and to evaluate the potential of MAGL inactivation as a therapeutic target for neurodegenerative diseases, it is crucial to assess how cell type-specific 2-AG metabolism influences brain homeostasis, particularly in relation to synaptic and neural circuit activities that are essential for cognitive function. This understanding is fundamental to evaluating the broader impact of 2-AG signaling on brain function, particularly cognition.

## 2. Results

### Inhibition of 2-AG metabolism in neurons impairs cognitive function

To explore cell type-specific 2-AG metabolism in brain function, we used *mgll* (the gene encoding MAGL) floxed mice by crossing them with specific Cre mice to generate total (tKO), astrocytic (aKO), and neuronal (nKO) MAGL knockout (KO) mice as described previously ^34^. Since 2-AG in microglial cells is primarily hydrolyzed by ABHD12 and MAGL does not play an important role in degrading 2-AG in these cells ^29,52^, we did not generate microglial MAGL KO mice for the present study. It has been previously demonstrated that 2-AG levels increased about 6 to 7-fold in tKO mice, 5-fold in nKO mice, and about 2-fold in aKO mice ^29,34^, suggesting that a large proportion of 2-AG in the brain is produced in neurons.

Given that the assessment of cognitive function is essential for evaluating therapies for neurodegenerative diseases, especially AD, and that the inhibition of 2-AG metabolism by inactivation of MAGL has been proposed as a therapeutic approach for neurodegenerative diseases like AD ^12,36,37,47–51^, we first assessed spatial learning and memory retention, which are critical cognitive functions. To this end, we used the Morris water maze (MWM) and novel object recognition (NOR) tests to evaluate learning and memory in these cell type-specific MAGL KO mice, as described previously ^34,37,40,53,54^. While there were no significant changes in behavioral performance of tKO mice in both MWM and NOR tests when compared to wild type (WT) mice, nKO mice displays significant impairments in learning and memory when compared to WT, tKO, and aKO mice (Figure 1A∼1F). In contrast, aKO mice showed enhanced learning in the MWM test (Figure 1A) when compared to WT mice. These data suggest that the inhibition of 2-AG metabolism in neurons is detrimental to cognitive function. The absence of impairments in learning and memory displayed in tKO mice suggests that cognitive deficits caused by the inactivation of neuronal MAGL are likely compensated for by the inactivation of MAGL in astrocytes.

**Figure 1.**
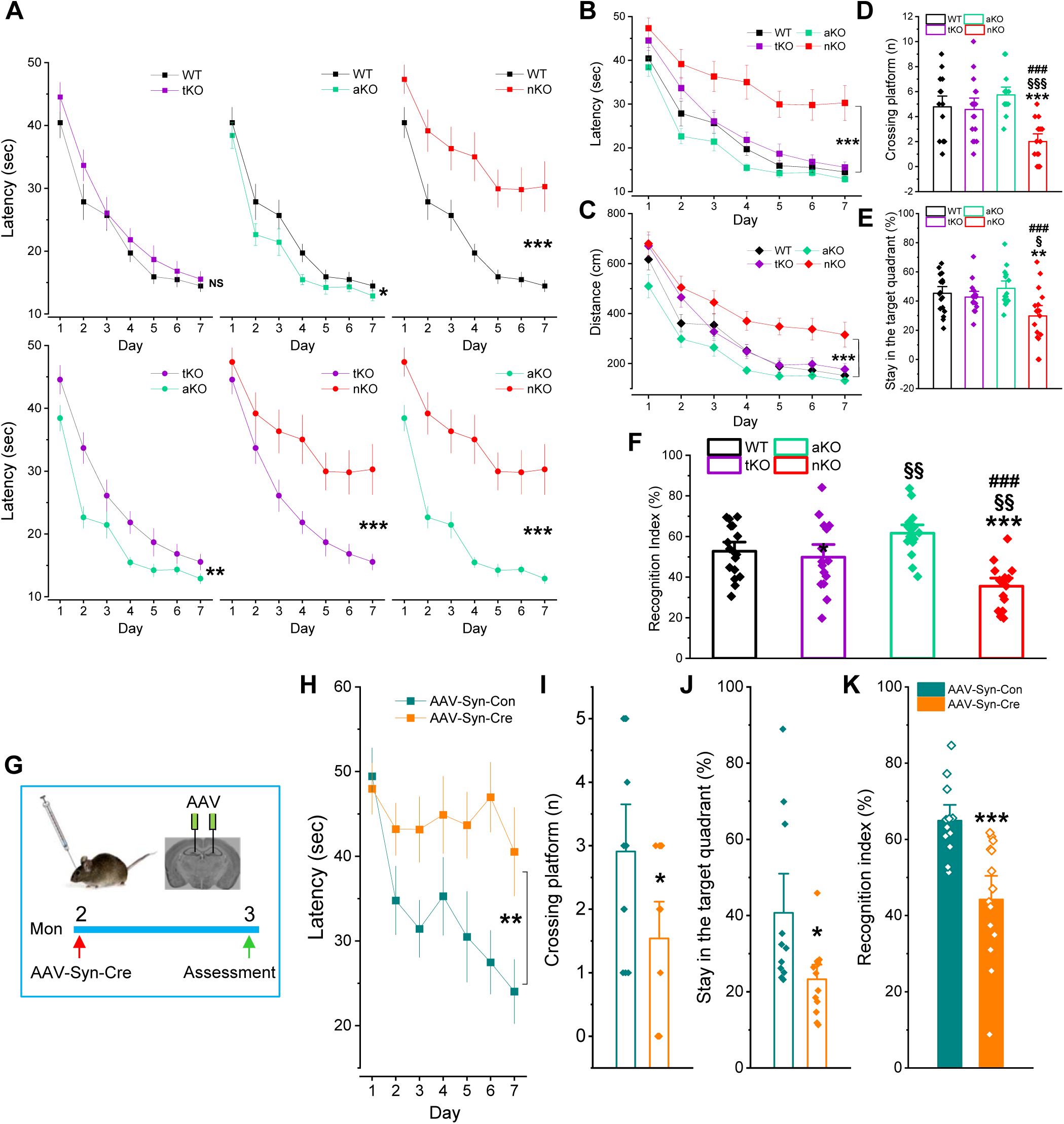
Inactivation of MAGL in neurons impairs learning and memory. (**A-C**) Spatial learning and memory were assessed using the Morris water maze (MWM) test in WT tKO, aKO, and nKO mice. The data are presented as means ±SEM. *P<0.05.**P<0.01. ***P<0.001 (ANOVA repeated measures, n=15∼18 animals/group). (**D-E**) The probe test was conducted 24 hours following 7 days of invisible platform training. The data are presented as means ±SEM. **P<0.01, ***P<0.001 compared with WT; §P<0.05, §§§P<0.001 compared with tKO; ###P<0.001 compared with aKO (ANOVA with Bonferroni post-hoc test, n=15∼18 animals/group). (**F**) Novel object recognition (NOR) test in WT, tKO, aKO, and nKO mice. The data are presented as means ±SEM (ANOVA with Bonferroni post-hoc test, n=16∼17 animals/group). (**G**) Schematic illustration of the experimental protocol. *Mgll* floxed mice at two months of age were stereotaxically injected with AAV-synapsin 1-cre (AAV-Syn-cre) vectors or AAV-Syn-control vectors into the hippocampus. The behavioral assessments were performed 30 days after injection of AAV vectors. (**H-J**) MWM test in *mgll* floxed mice injected with AAV-Syn-cre or control vectors. The data are means ±SEM. *P<0.05, **P<0.01 (ANOVA with Bonferroni post-hoc test, n=11∼13 animals/group). (**K**) NOR test in *mgll* floxed mice injected with AAV-Syn-cre or control vectors. The data are presented as means ±SEM. ***P<0.001 (ANOVA with Bonferroni post-hoc test, n=12∼14 animals/group).

The impaired cognitive function observed in nKO mice might result from genetic compensation during development. To test this possibility, we injected AAV-synapsin 1-cre (AAV-Syn-cre) vectors or AAV-Syn-control vectors into the hippocampus of *mgll* floxed mice at 2 months of age and assessed their behavioral performance 30 days after injection of AAV vectors (Figure 1G). As shown in Figure 1H∼1K, knockout of neuronal MAGL in the hippocampus led to deterioration in spatial learning and memory in both MWM and NOR tests. These results confirm that the inhibition of 2-AG breakdown in neurons impairs cognitive function.

### Inactivation of MAGL in neurons leads to deterioration in synaptic integrity

It is well recognized that 2-AG functions as a retrograde messenger modulating synaptic transmission and plasticity ^5,55^. Previous studies showed that both depolarization induced suppression of excitation (DSE) at cerebellar parallel fiber (PF) to Purkinje cell (PC) synapses and depolarization induced suppression of inhibition (DSI) at CA1 pyramidal neuron synapses are enhanced in tKO, aKO, and nKO mice ^29^. This suggests that MAGL in both astrocytes and neurons plays prominent roles in terminating 2-AG signaling at synaptic terminals ^29^. Since the structural integrity and functional properties of synapses are fundamental to cognitive function ^56–60^, and the hippocampal-entorhinal cortical networks are crucial for memory and cognitive functions, with the perforant path (PP)-dentate gyrus (DG) synapses serving as a major gateway into the hippocampus for information processing and encoding ^61–63^, we characterized spontaneous excitatory and inhibitory synaptic activities, input-output functions, and long-term potentiation (LTP) at this synapse. As shown in Figure 2A, while the frequency of spontaneous excitatory synaptic currents (sEPSCs) at hippocampal PP-DG synapses was reduced in cross tKO, aKO, and nKO mice, the amplitude of sEPSCs was reduced in both tKO and nKO mice, but not in aKO mice, with a more pronounced reduction in nKO mice. Similarly, tKO and nKO mice displayed a significant decrease in the input-output function, but not in aKO mice (Figure 2B). Importantly, LTP was impaired in nKO mice, but not in tKO mice (Figure 2C). In contrast, LTP was enhanced in aKO mice. The lack of LTP reduction in tKO mice may result from a compensatory effect of inactivating astrocytic MAGL.

**Figure 2.**
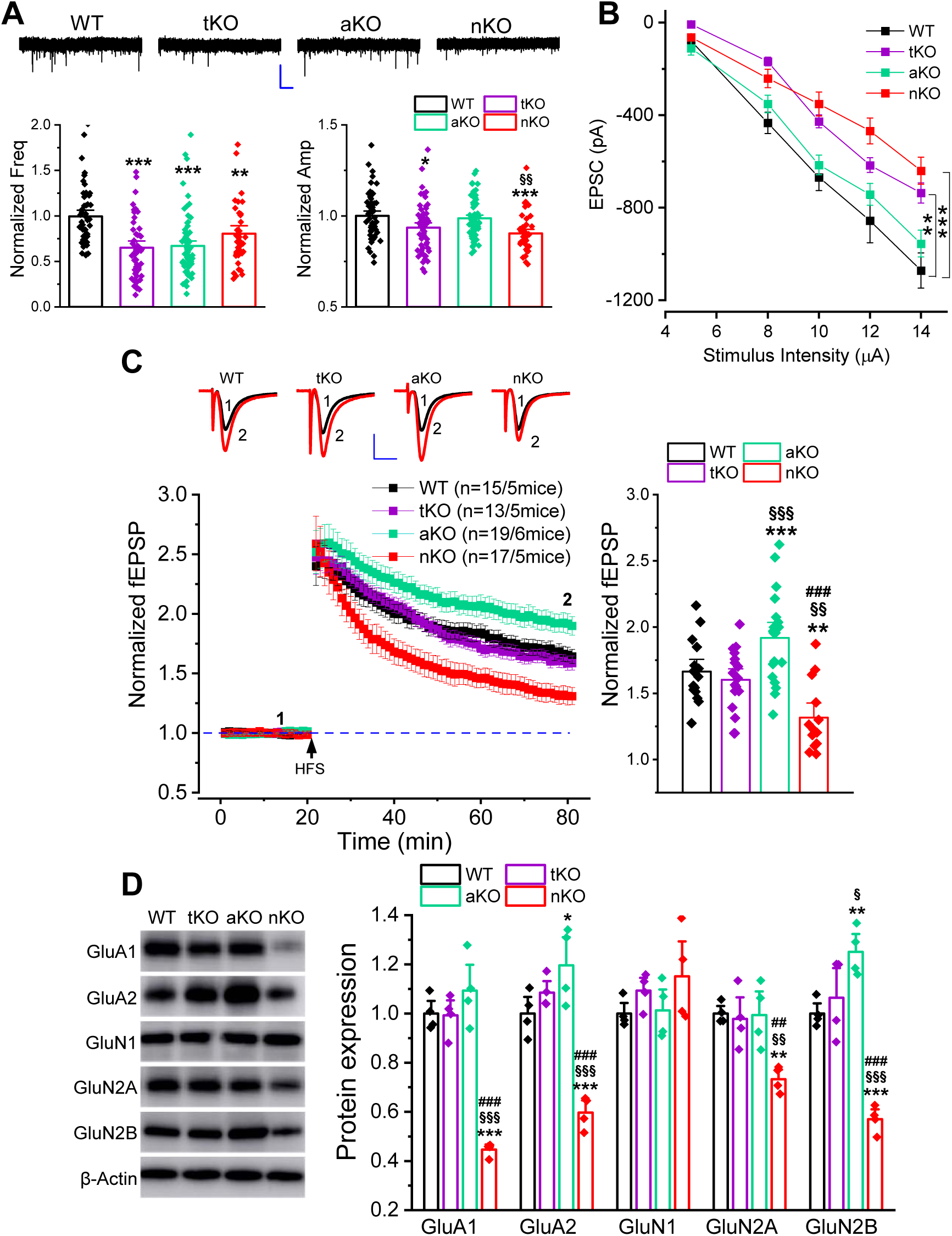
Selective loss of MAGL in neurons causes deterioration in synaptic integrity. (**A**) Spontaneous excitatory postsynaptic currents (sEPSCs) recorded at the perforant path (PP) synapses in the hippocampal dentate gyrus from WT, tKO, aKO and nKO mice. **P<0.01, ***P<0.001 compared with WT; §§P<0.01, compared with aKO (ANOVA with Bonferroni post-hoc test, n=6∼9 mice/group). (**B**) Input-output function of EPSCs at hippocampal PP synapses recorded from WT, tKO, aKO and nKO mice. **P<0.01, ***P<0.001 compared with WT (ANOVA with repeated measures, n=6∼9 mice/group). (**C**) Lon-term potentiation (LTP) recorded at hippocampal PP synapses from WT, tKO, aKO and nKO mice. Mean values of the potentiation of fEPSPs averaged from 56 to 60 min following high-frequency stimulation (HFS). The data are presented as means ±SEM. **P<0.01, ***P<0.001 compared with WT; §§P<0.01, §§§P<0.001 compared with tKO; ###P<0.001 compared with aKO (ANOVA with Bonferroni post-hoc test, n=5∼6 animals/group). (**D**) Immunoblot analysis of expression of glutamate receptor subunits in the hippocampus of WT, tKO, aKO and nKO mice. The data are means ±SEM. **P<0.01, ***P<0.001 compared with WT, §§P<0.01, §§§P<0.001 compared with tKO; ##P<0.01, ###P<0.001 compared with aKO (ANOVA with Fisher’s PLSD test post-hoc test, n=4 animals/group).

The reduced excitatory synaptic transmission and long-term synaptic plasticity observed in nKO mice are likely associated with decreases in the expression of glutamate receptors. Using immunoblot analysis, we found that the expression of AMPA glutamate receptor subunits GluA1 and GluA2, as well as NMDA receptor subunits GluN2A and GluN2B, was robustly downregulated in the hippocampus of nKO mice, but not in tKO mice (Figure 2D). The absence of impairments in LTP in tKO mice may result from compensatory mechanisms via inactivating astrocytic MAGL, as the expression of GluA2 and GluN2B was upregulated in aKO mice. This upregulation may also underlie the enhanced LTP in aKO mice.

To determine structural integrity of synapses in *mgll* KO mice, we assessed the morphology of hippocampal dendritic spines, as these structures are plastic and changes in the shape, size, and the density of dendritic spines are correlated with the strength of excitatory synaptic connections and memory formation ^64–69^. Using Golgi staining ^70^, we observed that the density of dendritic spines in hippocampal neurons was significantly reduced in nKO mice, but not in tKO mice (Figure 3A). In contrast, the density of dendritic spines was elevated in aKO mice. The reduction of hippocampal dendritic spine density in nKO mice indirectly indicates that the number of synapses might be reduced. To test this possibility, we used transmission electron microscopy (TEM) to directly detect synapses in the hippocampus. As shown in Figure 3B, the number of synapses in the hippocampus was increased in tKO and aKO mice, but significantly reduced in nKO mice, suggesting that inactivating neuronal MAGL results in a decrease in the number of synapses. Immunoblot data further confirm that the expression of synaptophysin (Syn) and postsynaptic density protein-95 (PSD-95), pre- and post-synaptic markers, was significantly decreased in the hippocampus of nKO mice, but not in tKO mice (Figure 3C). On the other hand, the expression of these synaptic markers was elevated in aKO mice. Additionally, the expression of ephrin type-B receptor 2 (ephB2) and sirtuin1 (sirt1), two important molecules regulating synaptogenesis, axon guidance, dendritic spine formation and synaptic maturation, expression and function of AMPA and NMDA glutamate receptors, long-term synaptic plasticity, and memory formation ^70–77^, was downregulated in nKO mice, but not in tKO mice. In contrast, their expression in aKO mice was upregulated (Figure 3C). These results indicate that the inactivation of MAGL in neurons leads to a deterioration in the structural and functional plasticity of synapses, which may underlie the impaired synaptic and cognitive functions observed in nKO mice. Concurrent inhibition of 2-AG metabolism in astrocytes can compensate for the disruption of structural and functional synaptic integrity caused by the knockout of neuronal MAGL.

**Figure 3.**
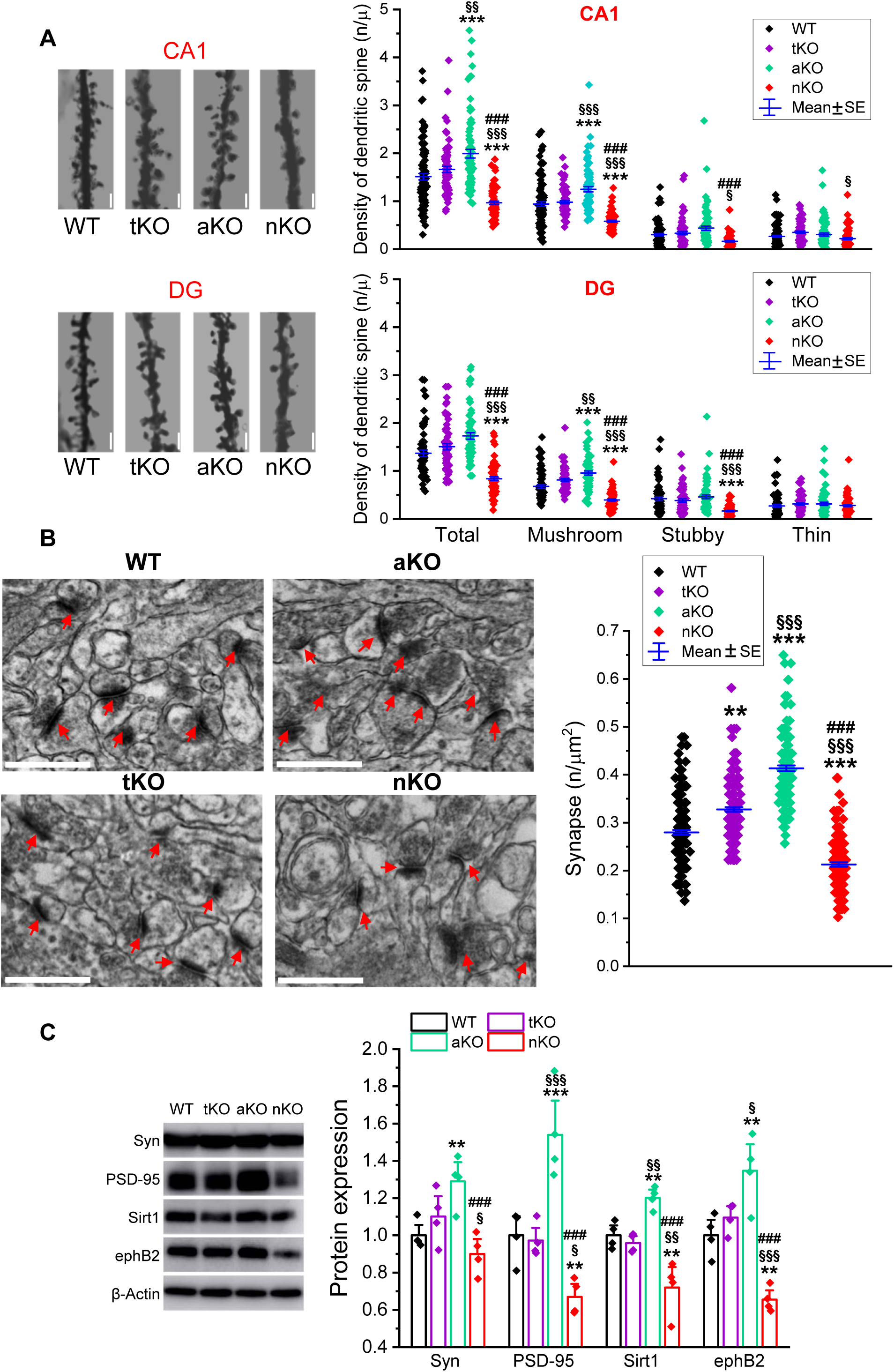
Inhibition of 2-AG degradation in neurons reduces synapses in the hippocampus. (**A**) Golgi staining of dendritic spines in hippocampal CA1 pyramidal neurons and dentate gyrus granule neurons from WT, tKO, aKO and nKO mice. ***P<0.001 compared with WT, §P<0.05, §§P<0.01, §§§P<0.001 compared with tKO; ###P<0.001 compared with aKO (ANOVA with Bonferroni post-hoc test group, n=5/group). Scale bars: 2 μm. (**B**) Transmission electron microscopy (TEM) images of hippocampal sections from WT, tKO, aKO and nKO mice. **P<0.01, ***P<0.001 compared with WT, §§§P<0.001 compared with tKO; ###P<0.001 compared with aKO (ANOVA with Bonferroni post-hoc test group, n=5/group). Scale bars: 800 nm. (**C**) Immunoblot analysis of the expression of synaptic proteins, including synaptophysin (Syn) and postsynaptic density protein 95 (PSD-95), and the proteins regulating synaptic structure and function, including sirtuin 1 (Sirt1) and ephrin type-B receptor 2 (ephB2) in the hippocampus of WT, tKO, nKO, and aKO mice. The data are means ±SEM. **P<0.01, ***P<0.001 compared with WT; §P<0.05, §§P<0.01, §§§P<0.001 compared with tKO; ###P<0.001 compared with aKO (ANOVA with Fisher’s PLSD post-hoc test, n=4 animals/group).

### Inhibition of 2-AG degradation in neurons disrupts intra-cortical circuit functional connectivity

Endocannabinoids have been shown to plays an important role in activity-dependent plasticity changes ^78,79^, which shape cortical circuit connectivity ^80,81^. To determine whether the impaired synaptic and cognitive functions in nKO mice are associated with changes in cortical circuit functional connectivity, we used laser scanning photostimulation (LSPS) combined with glutamate uncaging to map synaptic connectivity (Figure 4A), as described previously ^82–84^. Prefrontal layer 5 pyramidal neurons were targeted for patch clamp recording (Figure 4A) and LSPS/glutamate uncaging to probe both excitatory and inhibitory synaptic inputs (Figure 4B-D). After collecting multiple cells, pooled analysis revealed that excitatory synaptic inputs from layer 2/3 to layer 5 neurons are significantly reduced in nKO mice compared to WT controls (Figure 4E, G&H), but no deficit was observed in either tKO or aKO mice (Figure 4K, L, O &P). The inhibitory inputs onto the recorded layer 5 neurons, which primarily derive from both layer 2/3 and layer 5 locations, were also quantified. As shown in Figure 4F, I, J, M, N, Q&R, no significant changes in the overall strength of inhibitory synaptic inputs were observed in tKO, aKO, or nKO mice when compared with WT controls. These results suggest that the inhibition of 2-AG breakdown in neurons results not only in impairments in synaptic transmission and plasticity in the hippocampus but also leads to a disruption of cortical circuit functional connectivity. Importantly, concurrently enhancing 2-AG signaling in astrocytes may correct these abnormalities caused by the inactivation of neuronal MAGL, indicating a metabolic interplay of 2-AG signaling between astrocytes and neurons in maintaining brain homeostasis ^11,29^.

**Figure 4.**
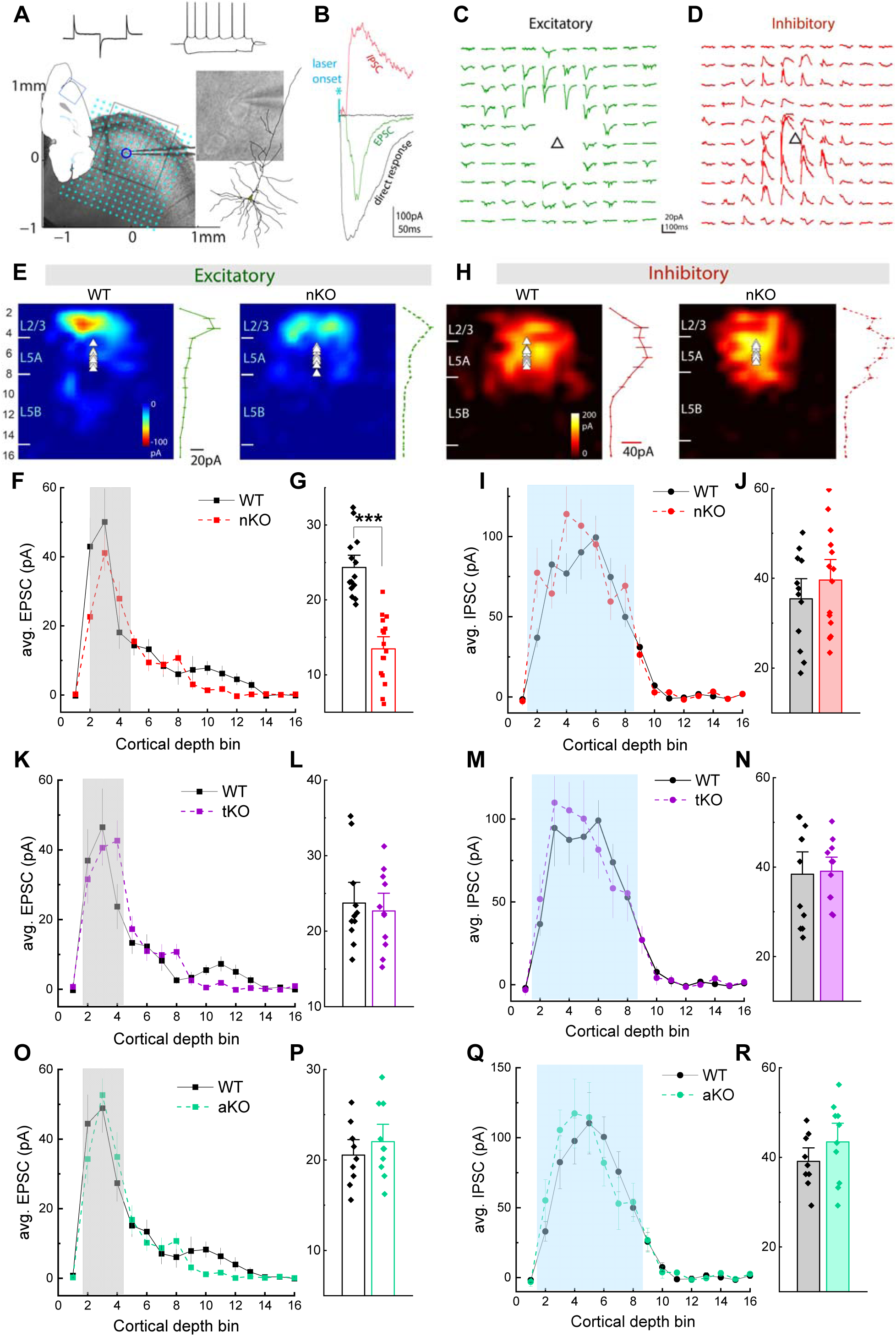
Inactivation of MAGL in neurons alters functional connectivity of cortical circuits. (**A**) Schematic illustration of a parasagittal prefrontal slice preparation, whole cell recording was made on a layer 5 pyramidal neuron. Sample tests of membrane/firing properties were listed above. Illustration of LSPS mapping, with a 16 x 16 stimulus grid overlaid, centered on the recorded layer 5 neuron, and aligned with pia surface. Cyan asterisks indicate uncaging locations. (**B**) LSLS mapping/glutamate uncaging at different locations (cyan asterisks) may elicit direct soma responses, excitatory synaptic currents (EPSC), or inhibitory synaptic currents (IPSC). Note their temporal relationship to laser onset. (**C∼D**) Representative mapping traces of a 10 × 10 grid of excitatory (**C**) and inhibitory (**D**) response arrays (boxed laser uncaging region in **A**). Triangles indicate soma position. (**E**) Averaged excitatory maps from L5 pyramidal neurons from WT and nKO mice. Line plot to the right denotes averaged responses from each cortical depth bin. (**F**) Averaged strength of excitatory synaptic inputs binned by cortical layers between WT and nKO neurons. (**G**) Comparison of pooled responses from L23 locations (box area in **F**), ***p < 0.001 (Student t test, n=14∼16/group). (**H**) Averaged inhibitory maps from L5 pyramidal neurons. (**I**) Averaged strength of inhibitory synaptic inputs binned by cortical layers. (**J**) No significant difference of averaged inhibitory inputs from combined layer 23 and layer 5 locations (p = 0.97, n=12∼14/group). (**K**) Averaged strength of excitatory synaptic inputs binned by cortical layers between WT and tKO neurons. (**L**) Comparison of averaged strength of excitatory synaptic inputs from combined L23 locations between WT and tKO neurons (p = 0.67, n=11/group). (**M**) Averaged inhibitory synaptic inputs to L5 neurons between WT and tKO neurons. (**N**) There was no significant difference for combined inhibitory inputs between WT and tKO neurons (p = 0.87, n=11/group). (**O**) Averaged excitatory maps of L5 pyramidal neurons from WT and aKO mice. (**P**) Comparison of averaged strength of excitatory synaptic inputs from combined L23 locations (p = 0.40, n=9∼10/group). (**Q**) Averaged inhibitory synaptic inputs to L5 neurons in WT and aKO neurons across cortical layers. (**R**) No significant difference was observed for combined inhibitory inputs between WT and aKO neurons (p = 0.23, n=9∼10/group).

### Enhancement of 2-AG in neurons induces widespread changes in the expression of genes associated with synaptic function

To further determine the mechanisms underlying the synaptic and cognitive deficits observed in nKO mice, we utilized 10x Genomics Chromium single-cell/nucleus RNA sequencing (sc/n-RNA-seq) technology to assess gene expression in the hippocampus of WT, tKO, aKO, and nKO mice. As shown in Figure S1A, there are 87 clusters of nuclei and cells in the Uniform Manifold Approximation and Projection (UMAP) plots. Figure 5A presents the UMAP plot with neurons and glial cells from WT, tKO, aKO, and nKO mice identified using gene markers, as shown in Figure S1B-J. We primarily analyzed the expression levels of genes in neurons, including excitatory and inhibitory neurons, and glial cells, including astrocytes and microglia, using the FindMarkers function in the Seurat package to compare gene expression in mgll KO mice with that in WT mice, as described previously ^16,34,85^. We found that the genetic deletion of MAGL leads to significant up- and down-regulation of differentially expressed genes (DEGs) in neurons and glial cells from tKO, aKO, and nKO mice compared to their WT counterparts (Figure S2A & B and Tables S1 & 2). Interestingly, the genetic inactivation of MAGL resulted in a higher number of upregulated DEGs in glial cells, while it caused a greater number of downregulated DEGs in neurons (Figure S2A & B), highlighting a fascinating divergence in gene expression responses between these cell types. Gene Ontology (GO) analysis of DEGs in neurons and glial cells using the clusterProfiler R package reveals that neurons and glial cells showed significant dysregulation of GO pathways related to synapse organization, axon guidance, cellular respiration and mRNA processing across these transgenic mice (Table S3 and 4).

**Figure 5.**
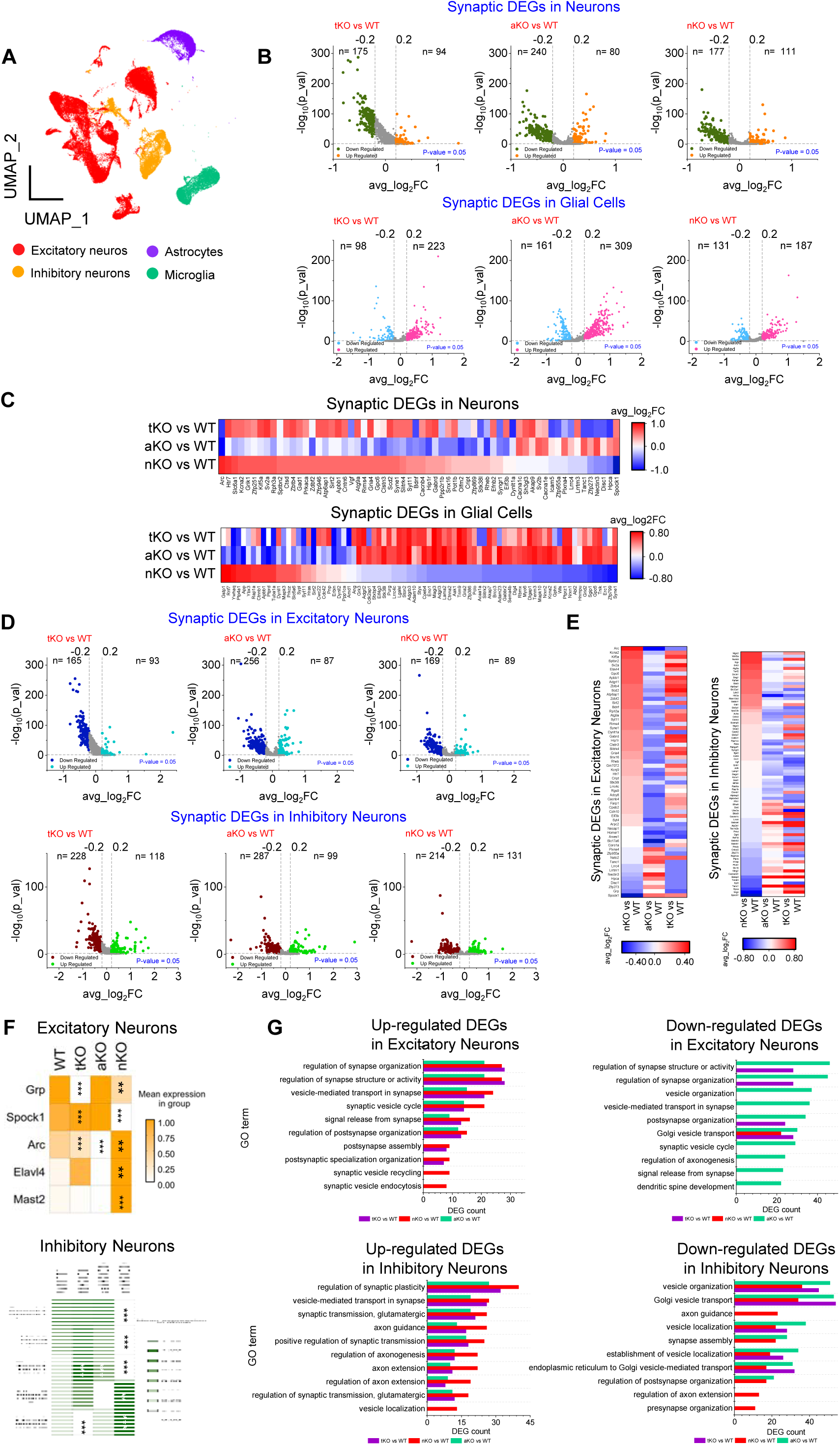
Inhibition of 2-AG metabolism alters expression of synaptic genes in hippocampal neurons. (**A**) UMAP visualization of integrated cells from WT, tKO, aKO, and nKO mice. (**B**) Volcano plots of differentially expressed genes (DEGs) in neurons (Upper) and glial cells (Lower) from tKO, aKO, and nKO mice compared with WT. (**C**) Heatmaps representing the fold change (avg_log2FC) of some DEGs which are related to synaptic function in neurons and glial cells from tKO, aKO, and nKO mice. (**D**) Volcano plots display the number of DEGs in excitatory and inhibitory neurons. (**E**) Heatmaps showing avg_log2FC of several synaptic DEGs in hippocampal excitatory and inhibitory neurons from tKO, aKO and nKO mice. (**F**) Heatmaps showing representative synaptic DEGs in excitatory and inhibitory neurons. ***P<0.001 compared with WT. (**G**) Bar plots depicting Gene Ontology (GO) term enrichment for synaptic function-related enriched pathways in excitatory and inhibitory neurons from hippocampus across tKO, aKO and nKO mice.

To explore whether impaired synaptic and cognitive functions in nKO mice are associated with changes in gene expression, we specifically analyzed synaptic DEGs in neurons and glial cells across tKO, aKO, and nKO groups (Figure 5B). In nKO mice, 1,080 synaptic genes were identified in neurons and 288 of these genes were significantly differentially expressed (Table S5). The expression of synaptic DEGs in neurons shows opposite trends between nKO and aKO mice. The heatmaps in Figure 5C (Top) list 61 interesting synaptic DEGs in neurons with opposite trends between the nKO and aKO groups. Among these DEGs, upregulated synaptic DEGs are more prevalent than downregulated DEGs in hippocampal neurons from nKO mice. Additionally, hippocampal neurons from tKO mice display a similar expression pattern of these synaptic DEGs to those from nKO mice. For example, *hpca* and *disc1*, which regulate long-term synaptic plasticity and NMDA receptor dynamics ^86,87^, exhibit similar trends between nKO and tKO mice, but opposite trends to those in neurons from aKO mice. This likely results from neuronal MAGL being inactivated in both tKO and nKO mice. Notably, *spock1*, *arc*, *kcna2*, *grik1*, and *htr7*, which are known regulators of the structural and functional plasticity of synapses ^88–94^, showed robustly altered expressions in neurons from nKO mice compared to WT mice (Figure S3A).

Similar trends in expression of synaptic DEGs were also observed in glial cells resulting from the inactivation of MAGL (Table S6). We identified 318 significantly changed synaptic DEGs, with 79 of these showing opposite trends between nKO mice and aKO mice (Figure 5C bottom). Surprisingly, glial cells from tKO mice exhibit an expression pattern of these synaptic DEGs that is consistent with the pattern observed in aKO mice, but not nKO mice. For instance, *syne1* and *tnik*, which regulate glutamate receptor expression and signaling ^95,96^, show comparable trends between aKO and tKO mice but display opposite trends in glial cells from nKO mice. Among synaptic DEGs in glial cells, *hnrnpu*, *abl2*, *gstp1*, *ptprz1*, and *rnf7* ^97–101^, which are involved in regulation of synaptic transmission, synapse localization, neuritogenesis, glutamatergic synaptic activity, are dysregulated DEGs (Figure S3B). These altered synaptic DEGs in glial cells resulting from the inactivation of MAGL may contribute to the structural and functional plasticity of synapses.

To further investigate synaptic DEGs in subtypes of neurons, we classified these synaptic DEGs in excitatory and inhibitory neurons and found that cell type specific inhibition of 2-AG metabolism led to diverse transcriptomic profile (Figure S2C and 2D, Table S7 and 8), affecting synaptic function, translation function, and oxidative respiration (Table S9 and 10). Synaptic genes in excitatory and inhibitory neurons across *mgll* genotypes were detected, as shown in Tables S11 and S12. Figure 5D illustrates significantly altered synaptic genes in both excitatory and inhibitory neurons. Additionally, we identified 60 crucial synaptic DEGs in excitatory neurons and 346 in inhibitory neurons from nKO mice, with expression trends opposite to those in aKO mice (Figure 5E). Figure 5F lists the most significant synaptic DEGs, including *grp*, *spock1*, *arc*, *elavl4*, *hpca*, *tanc1*, and *nlgn2*, which regulate synaptic structure and function ^87,90,102–108^ across different genotypes. Several synaptic DEGs are altered in both excitatory and inhibitory neurons from all *mgll* KO mice, but some DEG changes display cell type specific. For instance, *elavl4* is only changed in excitatory neurons, while *nlgn2* is altered in inhibitory neurons (Figure 5F). The GO term analysis highlights that synaptic functions in both excitatory neurons are significantly impacted across all genotypes. However, certain GO terms show specificity to different genotypes (Figure 5G, top). For instance, the upregulation of “synaptic vesicle recycling” is uniquely observed in nKO mice. Similarly, specific trends are noted in inhibitory neurons (Figure 5G, bottom), indicating genotype-specific effects. For example, the downregulated “axon guidance” term is specific to nKO mice. These results suggest that while the knockout of MAGL broadly affects various aspects of synaptic functions, certain processes are distinctly altered depending on the genotype.

In glial cells, including astrocytes and microglia, inactivation of MAGL induced significant enrichment in DEGs associated with various biological processes essential to autophagy, Golgi vesicle transport, regulation of cellular macromolecule biosynthetic processes, and positive regulation of proteolysis (Tables S15 and S16) among tKO, aKO, and nKO mice. Importantly, inactivation of MAGL led to changes in numbers of synaptic DEGs across genotypes, as detailed in Tables S13 and S14 and shown in Figure 6A. We detected a significant number of synaptic genes that are up- or down-regulated in astrocytes and microglia across genotypes (Figure S2E and 2F, Table S17 and 18). Notably, 53 synaptic DEGs in astrocytes and 25 synaptic DEGs in microglia exhibited opposite expression trends between aKO and nKO mice (Figure 6B). GO analysis indicates that synaptic DEGs in astrocytes are involved in regulating various components of synaptic activities, with some being specific to nKO type, such as “regulation of axonogenesis” and “vesicle organization” (Figure 6C). Synaptic DEGs in microglia also display a similar patten in nKO mice (Figure 6D). Among the GO terms regulating synaptic functions, more terms are specific to nKO. Notably, *kcna2*, *adam23*, *il33, ank2,* and *socs2* in astrocytes, as well as *hnrnpu*, *dynll1, tsc2, egr3,* and *ptgds* in microglia, which are involved in the regulation of synaptic functions including synaptic transmission and plasticity, neurite outgrowth, axonal plasticity, axonal trafficking and formation of synapse ^89,98,109–117^, were significant altered in nKO mice (Figure 6E). These findings suggest that inhibition of 2-AG metabolism results in the reversal of expression of many important synaptic DEGs in both astrocytes and microglia between nKO and aKO mice.

**Figure 6.**
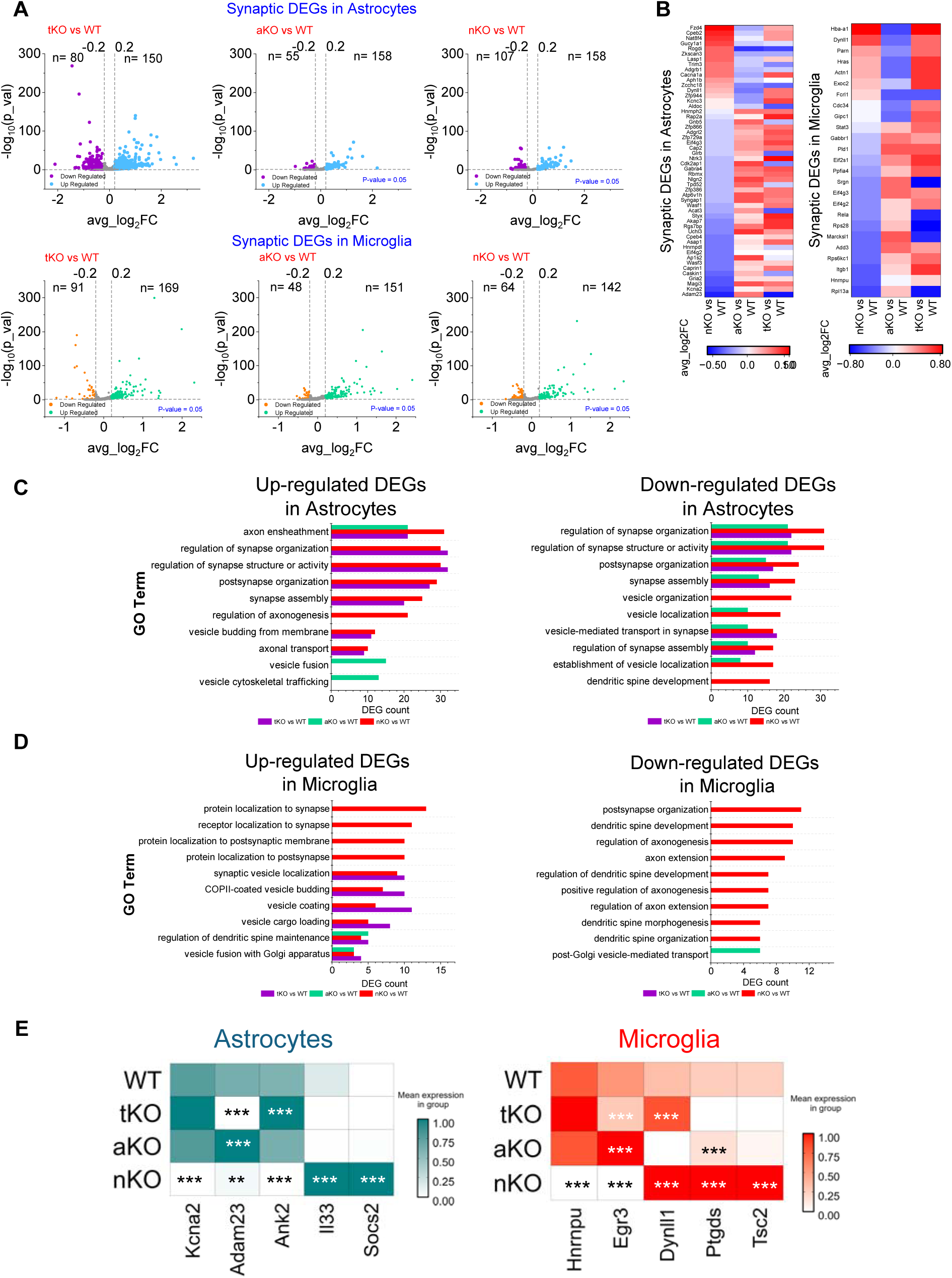
Inactivation of MAGL leads to changes in synaptic gene expression in microglia and astrocytes. (**A**) Volcano plots displaying the numbers of DEGs in astrocytes and microglia across tKO, aKO and nKO groups. (**B**) Heatmaps display genes crucial for synaptic function in astrocytes and microglia. (**C∼ D**) Histogram plots showing the numbers of synaptic DEGs in different GO terms related to synaptic structure and function. (**E**) Heatmaps illustrating representative DEGs in astrocytes and microglia. ***P <0.001 compared with WT.

### Inactivation of neuronal MAGL impairs adult neurogenesis

Adult-born hippocampal neurons are important for memory and cognitive plasticity ^118–120^. We found that inactivation of MAGL caused changes in the expression levels of DEGs related to adult neurogenesis in neurons and glial cells (Figure 7A and Table S22∼24). The heatmaps in Figure 7A show that DEGs related to adult neurogenesis in nKO mice display opposite trends compared to those in aKO mice. For example, the expression of *gsn*, *slc12a2*, and *app*, which control adult neurogenesis in the hippocampus under different conditions ^121–123^, was increased in glial cells from nKO mice, while their expression levels in aKO mice were significantly decreased or showed a downregulated trend (Figure 7B). This suggests that inactivation of neuronal MAGL might impair adult neurogenesis by altering DEGs in glial cells and neurons. Indeed, we observed that bromodeoxyuridine or 5-bromo-2’-deoxyuridine (BrdU)- and doublecortin (DCX)-positive cells in the hippocampal dentate gyrus region are significantly reduced in nKO mice when compared with WT, tKO, and aKO mice (Figure 7C&D). These results suggest that inhibition of 2-AG degradation in neurons affects adult neurogenesis, which may contribute to the deterioration in synaptic and cognitive functions observed in nKO mice.

**Figure 7.**
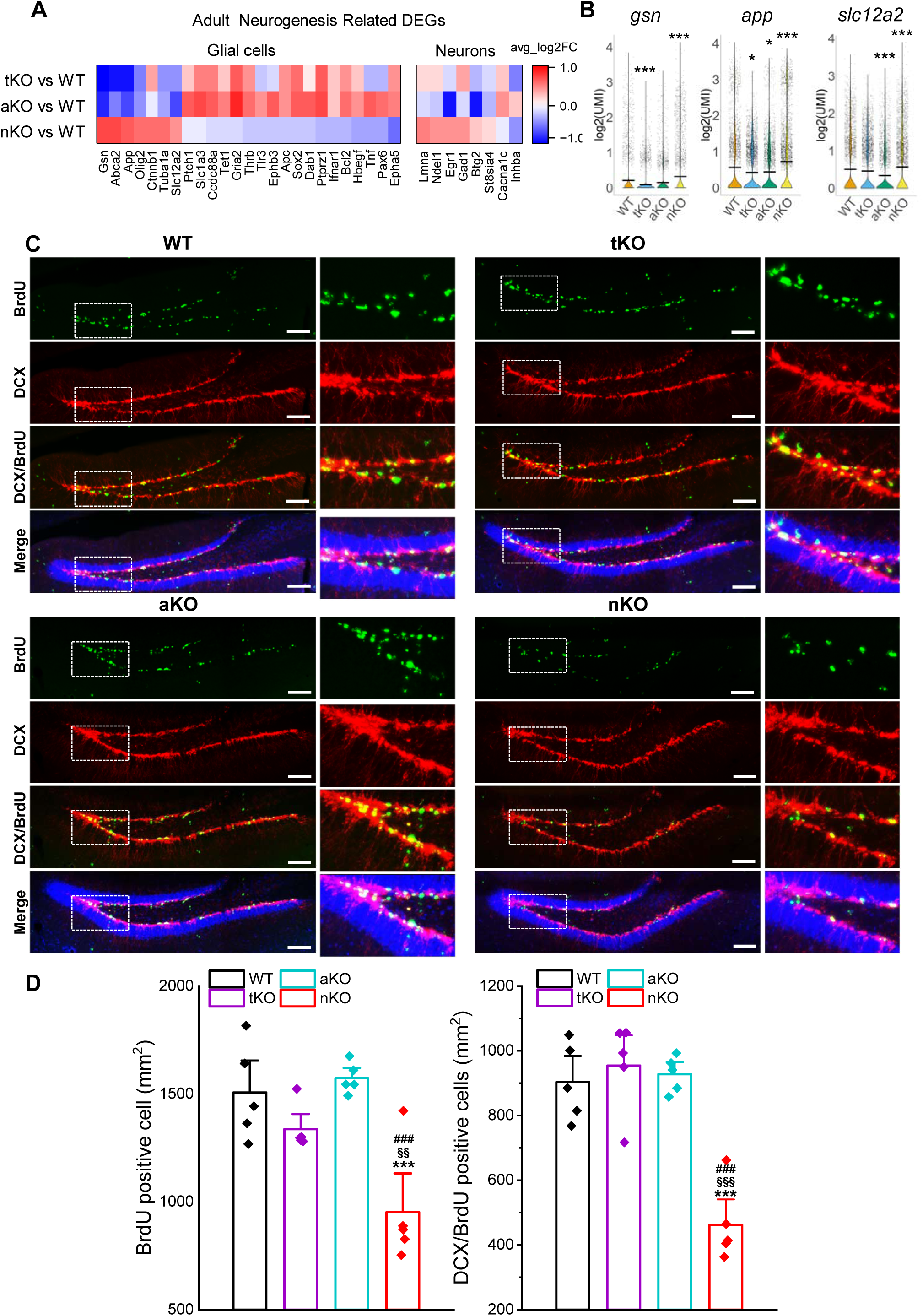
Inhibition of 2-AG metabolism in neurons impairs adult neurogenesis. (**A**) Heatmaps showing multiple DEGs related to adult neurogenesis in glial cells and neurons from tKO, aKO and nKO mice. Notably, the trends of these DEGs are opposite between aKO and nKO mice. (**B**) Representative violin plots of DEGs related to adult neurogenesis. *p<0.05, **p<0.01, ***p<0.001 compared with WT. (**C**) Immunostaining images of BrdU (green), DCX (red), and DAPI (blue) in the dentate gyrus region of the hippocampus across different genotypes. (**D**) Quantitative analysis of the number of BrdU and DCX/BrdU double-positive cells in the hippocampus from WT, tKO, aKO, and nKO mice. Data are represented as means ± SEM. ***p < 0.001 compared with WT; §§P<0.01, §§§P<0.001 compared with tKO; ###P<0.001 compared with aKO (ANOVA with Bonferroni post-hoc test, n=5 animals/group). Scale bars: 50 um.

## 3. Discussion

It is well recognized that 2-AG acts as a retrograde messenger modulating synaptic transmission and plasticity at both inhibitory GABAergic and excitatory glutamatergic synapses in the brain ^1–8^. In particular, 2-AG displays anti-inflammatory and neuroprotective properties in response to various stimuli or harmful insults, thereby maintaining brain homeostasis ^10–18,23^. However, 2-AG is rapidly degraded by several enzymes following its synthesis ^12,24^, with MAGL being the key enzyme responsible for degrading 2-AG in the brain ^25–29^. It has been shown that pharmacological or genetic inactivation of MAGL reduces neuropathology and prevents synaptic and cognitive declines in animal models of neurodegenerative diseases ^12,34,37–46^. Consequently, inhibit 2-AG metabolism by inactivating MAGL has been proposed as a potential therapeutic strategy for neurodegenerative diseases, including AD ^12,36,37,47–51,124^. Correspondingly, many pharmacological MAGL inhibitors have been developed or are currently in development ^125,126^.

Despite this, the impact of cell type-specific 2-AG metabolism on synaptic structure and function, and cognitive abilities has not been thoroughly evaluated, which is particularly important if MGAL inactivation is to be considered as a therapy for AD. To address this, we utilized genetically engineered *mgll* floxed mice ^34^, to assess synaptic and cognitive functions in total, astrocytic, and neuronal MAGL knockout mice. In the present study, we observed that inactivating MAGL in neurons impairs learning and memory in mice. This cognitive abnormity appears to result from decreases in the expression of synaptic proteins, the number of synapses, long-term synaptic plasticity, cortical circuit functional connectivity, and neurogenesis. Interestingly, the synaptic and cognitive deficits induced by neuronal MAGL inactivation could be rescued by deleting astrocytic MAGL. Transcriptomic analyses at the single-cell/nucleus levels revealed that inactivation of MAGL in neurons leads to widespread changes in the expression of genes associated with synaptic function. These findings suggest that there is crosstalk in 2-AG signaling between astrocytes and neurons in maintaining brain homeostasis, and that excessive 2-AG in neurons alone is detrimental to cognitive function.

It has been observed that the activity or expression MAGL is altered in patients with AD ^127–129^ and in experimental models of TBI ^34,130^, which may contribute to the disruption of brain homeostasis, thereby leading to brain disorders. For instance, AAV vector-mediated overexpression of MAGL in hippocampal glutamatergic neurons, which decreases in 2-AG in excitatory neurons, results in impaired short-term synaptic plasticity of excitatory synapses and increased anxiety-like behavior ^131^. In the present study, we found that increasing 2-AG levels in neurons through selective knockout of MAGL induces profound synaptic and cognitive deterioration in mice. This suggests that maintaining proper 2-AG metabolism in neurons is essential for normal synaptic and cognitive functions. However, we did not observe significant synaptic and cognitive impairments in global MAGL knockout mice (tKO), even though neuronal MAGL is also inactivated in these tKO mice. This lack of impairment is likely due to the concurrent elevation of 2-AG levels in astrocytes in the tKO mice. Our results indicate that increasing 2-AG levels in astrocytes enhances the expression of synaptic proteins, increases the number of synapses, and improves learning and memory. These changes, driven by augmented 2-AG signaling in astrocytes, appear to compensate for the impairments caused by neuronal MAGL inactivation, suggesting a crosstalk between neurons and astrocytes in maintaining synaptic and cognitive functions as well as brain homeostasis. This is supported by a previous study that observed transcellular shuttling of 2-AG and related metabolites between neurons and astrocytes as a mechanism by which these cells coordinately regulate endocannabinoid-eicosanoid pathways in the nervous system ^29^. Furthermore, a recent study strengthens the evidence for neuron-astrocyte interplay in 2-AG signaling by showing that global knockout of MAGL protects the brain against TBI, whereas neuronal MAGL knockout mice do not display this protection ^34^. Transcriptomic analyses at the single-cell/nucleus levels support this hypothesis. We found that inactivation of neuronal MAGL induces changes in the expression of genes associated with synaptic function that are opposite in astrocytes. These opposing changes in gene expression between neurons and glial cells may underlie the compensatory effects observed when neuronal MAGL inactivation-induced synaptic and cognitive deficits are counteracted by inhibiting 2-AG metabolism in astrocytes, thereby preserving synaptic and cognitive functions.

Our results suggest that crosstalk in 2-AG signaling between astrocytes and neurons is crucial for maintaining synaptic and cognitive functions, and that excessive 2-AG in neurons alone is detrimental to cognitive function. This understanding is essential for evaluating the broader impact of 2-AG signaling on brain function, particularly cognition. For instance, MAGL has emerged as an attractive therapeutic target for neurodegenerative diseases, with many MAGL inhibitors either already developed or in development. However, based on our findings, global inactivation of MAGL (*e.g.*, through pharmacotherapies) may not be the optimal approach for achieving ideal therapeutic outcomes in neurodegenerative diseases. Instead, selectively targeting MAGL in astrocytes to enhance 2-AG signaling could offer a more effective therapeutic strategy for neurological disorders ^12^.

## 4. Experimental Section Animals

Mgll^flox/flox^ animals were generated by the Texas A&M Institute for Genomic Medicine, as described previously ^34^. Briefly, the mutant allele carries the LoxP sites flanking Exon 2 of the gene. The LoxP sites were introduced by homologous recombination with a targeting vector in the C57BL/6N ES cell line JM8. The TIGM proprietary vector carried Neomycin transferase cassette for selection of correctly targeted clones; this cassette was flanked by Frt recombination sites that were later removed by breeding with the “Flpe deleter” mouse line to produce conditional-ready knockout (Mgll^flox/flox^). Deletion of the targeted exons was confirmed by crossing Mgll^flox/flox^ mice with Tg(Sox2-cre)1Amc/J (JAX Stock No: 004783) Sox2-Cre, resulting in a total/global MAGL knockout. Correct targeting and recombination were confirmed by Long Distance PCR and sequencing. Neuronal and astrocytic MAGL KO mice (nKO and aKO) were generated by crossing *mgll* ^flox/flox^ mice with Syn1-cre mice (JAX Stock No: 003966) and GFAP-cre mice (JAX Stock No: 024098), respectively. Animals were randomly assigned to groups from different genotypes. The number of animals per experimental group was calculated through a power analysis, using a power of 80%, an α of 0.05, and variables based on our previously published results from similar experiments. Both male and female mice at ages of 8 to 12 weeks were used in the present study.

All animal studies were performed in compliance with the US Department of Health and Human Services Guide for the Care and Use of Laboratory Animals, and the care and use of the animals reported in this study were approved by the Institutional Animal Care and Use Committee of University of Texas Health Science Center at San Antonio.

### Western blots

Western blot assay was conducted to determine expression of glutamate receptor subunits GluA1, GluA2, GluN1, GluN2A, and GluN2B, synaptophysin (Syn), PSD-95, ephrin type-B receptor 2 (EphB2), and sirtuin 1 (Sirt1) in hippocampal tissues from WT and MAGL KO mice, including tKO, nKO, and aKO mice. Hippocampal tissue was extracted and immediately homogenized in RIPA lysis buffer and protease inhibitors, and incubated on ice for 30 min, then centrifuged for 10 min at 10,000 rpm at 4°C. Supernatants were fractionated on 4-15% SDS-PAGE gels (Bio-Rad) and transferred onto PVDF membranes (Bio-Rad). The antibodies used to detect the expression of proteins are listed in Key resources table. The membrane was incubated with specific antibodies at 4°C overnight. The blots were washed and incubated with a secondary antibody (goat anti-rabbit 1:2,000, Cell Signaling) at room temperature for 1 hr. Proteins were visualized by enhanced ^40,41^ chemiluminescence (ECL, Amersham Biosciences, UK). The densities of specific bands were quantified by densitometry using GE/Amersham Imager 680 UV. Band densities were normalized to the total amount of protein loaded in each well as determined by mouse anti β-actin (Santa Cruz), as described previously ^18,37,40,41,53^.

| REAGENT or RESOURCE | SOURCE | IDENTIFIER |
| --- | --- | --- |
| **Antibodies** |  |  |
| Rabbit polyclonal anti-GluA1 | Abcam | Cat# ab31232; RRID: AB_2113447 |
| Rabbit monoclonal anti-GluA2 | Abcam | Cat# ab133477; RRID: AB_2620181 |
| Mouse monoclonal anti-GluN1 | Abcam | Cat# ab109182; RRID: AB_10862307 |
| Rabbit polyclonal anti-GluN2A | Millipore | Cat# 07-632; RRID: AB_310837 |
| Rabbit polyclonal anti-GluN2B | Abcam | Cat# ab65783; RRID: AB_1658870 |
| Mouse monoclonal anti-PSD95 | Abcam | Cat# ab99009; RRID: AB_10676078 |
| Mouse monoclonal anti-PSD95 | Abcam | Cat# ab2723; RRID: AB_303248 |
| Mouse monoclonal anti-Synaptophysin | Abcam | Cat# ab8049; RRID: AB_2198854 |
| Rabbit polyclonal anti-EPHB2 | Abcam | Cat# ab5418; RRID: AB_2262360 |
| Mouse monoclonal anti-SIRT1 | Abcam | Cat# ab110304; RRID: AB_10864359 |
| Goat anti-rabbit secondary antibody | Cell Signaling | Cat# 7074; RRID: AB_2099233 |
| Mouse monoclonal anti-BrdU antibody | Sigma | Cat# B8434; RRID:AB_476811 |
| Rabbit polyclonal anti-DCX antibody | Abcam | Cat# ab18723; RRID:AB_732011 |
| Mouse monoclonal anti-β-Actin | Santa Cruz | Cat# sc-47778; RRID: AB_626632 |

### Hippocampal slice preparation

Hippocampal slices were prepared from mice as described previously ^37,40,53,70^. Briefly, after decapitation, brains were rapidly removed and placed in cold oxygenated (95% O_2_, 5% CO_2_) artificial cerebrospinal fluid (ACSF) containing: 125.0 NaCl, 2.5 KCl, 1.0 MgCl_2_, 25.0 NaHCO_3_, 1.25 NaH_2_PO_4_, 2.0 CaCl_2_, 25.0 glucose, 3 pyruvic acid, and 1 ascorbic acid. Slices were cut at a thickness of 350-400 µm and transferred to a holding chamber in an incubator containing ACSF at 36 °C for 0.5 to 1 hour and maintained in an incubator containing oxygenated ACSF at room temperature (∼22-24 °C) for >1.5 h before recordings. Slices were then transferred to a recording chamber where they were continuously perfused with 95% O_2_, 5% CO_2_-saturated standard ACSF at ∼32-34 °C.

### Electrophysiological recordings

Whole-cell patch-clamp recordings were made using an Axopatch-200B patch-clamp amplifier (Molecular Devices, Sunnyvale, CA) under voltage clamp as described previously ^37,132^. Pipettes (2-4 MΩ) were pulled from borosilicate glass with a micropipette puller (Sutter Instrument, Novato, CA). The internal pipette solution contained (in mM): 90.0 CsCH_3_SO_3_, 40.0 CsCl, 10.0 HEPES, 5.0 CaCl_2_, 4.0 Mg_2_ATP, 0.3 Na_2_GTP, and 5.0 QX-314, or 130 KCH_3_SO_4_, 10 KCl, 4 NaCl 10 HEPES, 0.1 EGTA, 4 Mg_2_ATP, 0.3 Na_2_GTP, and 5 QX-314. The membrane potential was held at -70 mV. Evoked excitatory postsynaptic currents (EPSCs) in dentate granule neurons were recorded in response to stimuli of perforant path synapses (PP) at a frequency of 0.05 Hz using bipolar tungsten electrodes, as described previously ^70,133^. Spontaneous EPSCs were recorded, and the input-output function at PP-DG synapses was established by increasing the stimulus intensity. SR95531 (gabazine, 1 µM) was used to block GABAergic synaptic transmission during recordings of sEPSCs, and DL-AP5 (50 µM) and CNQX (20 µM) were included in the bath solution to block AMPA and NMDA currents during recordings of sIPSCs. The amplitude, frequency, and kinetics of sIPSCs/sEPSCs were analyzed using the MiniAnalysis program.

Field EPSP (fEPSP) recordings at hippocampal Schaffer-collateral synapses in response to stimuli at a frequency of 0.05 Hz were made using an Axoclamp-2B patch-clamp amplifier in bridge mode, as described previously ^37,40,41,53^. Recording pipettes were pulled from borosilicate glass with a micropipette puller (Sutter Instrument), filled with artificial ACSF (∼4 MΩ). As described previously ^37,40,41,53,70^, long-term potentiation (LTP) at perforant path-dentate gyrus synapses was induced by a high-frequency stimulation (HFS) consisting of three trains of 100Hz stimulation (1 sec duration and a 20 sec inter-train interval).

### Laser scanning photostimulation (LSPS) combined with glutamate uncaging to map prefrontal circuit connectivity

To investigate whether genetic inactivation of MAGL affects cortical circuit function and connectivity, we used laser scanning photostimulation (LSPS) combined with glutamate uncaging to map synaptic connectivity onto the L5 excitatory pyramidal neurons in the prefrontal cortex, as described previously ^84,134^. nKO, tKO, and aKO mice, along with their WT littermates, were sacrificed, and 300 μm parasagittal brain slices were prepared ^83^. The slices were perfused in modified ACSF with elevated calcium and magnesium, saturated with 95% O_2_ and 5% CO_2_, and containing (in mM) 126 NaCl, 2.5 KCl, 26 NaHCO_3_, 4 CaCl_2_, 4 MgCl_2_, 1.25 NaH_2_PO_4_, and 10 glucose. The ACSF also included 0.2 mM MNI-caged glutamate and 5 μM R-CPP to block NMDA receptors and prevent short-term plasticity changes. L5 neurons with pyramidal-shaped soma were targeted for whole-cell patch clamp recordings. To minimize truncation of dendritic structure and preserve connectivity, we selected neurons with soma located > 50 μm below the slice surface.

LSPS mapping was performed in a customized recording chamber mounted on a motorized stage (MPC-78, Sutter Instruments). We used a microscope (Olympus BX51WI) equipped with a 4× objective lens (NA 0.16; Olympus) and a 60× water immersion objective (NA 0.9, Olympus). After neurons were targeted and whole-cell configuration was obtained, membrane properties were tested by a +5-mV voltage step under voltage-clamp. Neurons were then injected with a series of 1-second current steps (-100 pA to 500 pA in 50 pA increments) in current-clamp mode to test their firing properties. The electrode internal solution contained (in mM): 130 K-gluconate, 10 HEPES, 4 KCl, 4 ATP-Mg, 2 NaCl, 0.3 GTP-Na, 1 EGTA, and 14 phosphocreatine (pH 7.2, 295-300 mOsm). After collecting neuronal firing properties, we carefully switched to the 4× objective without disrupting the seal for LSPS mapping. One-millisecond, 20-mW UV laser pulses (355 nm; DPSS Lasers) were scanned onto slices in a 16×16 stimulation grid with 100 μm spacing. The stimulation grid was left-right centered on the targeted neuron, and the top row was aligned with the pia surface. Neurons were first voltage clamped at -70 mV (resting membrane potential) to collect their excitatory input maps, followed by 0 mV (reversal potential for AMPA receptors) to collect inhibitory inputs. Neuronal signals were conditioned with a Multiclamp 700B amplifier (Molecular Devices), digitized at 20 kHz, and acquired using BNC-6259 boards (National Instruments, Austin, TX). Laser power and timing were controlled by an optic shutter (Conoptics, model 3050) and a mechanical shutter (Uniblitz VCM-D1). Digital images were acquired using a CCD camera (Retiga 2000DC, QImaging, Canada). Data acquisition and analyses were performed using Ephus software ^134^.

### Golgi–Cox staining

Golgi–Cox staining was used to detect dendritic spines of hippocampal neurons as we described previously with modification ^37,53,70,135^. WT or MAGL KO mice were anesthetized with ketamine/Xylazine (200/10 mg/kg) and subsequently transcardially perfused with ice-cooled saline for 5 min. The brain was dissected out and processed with Golgi-Cox Impregnation & Staining System according to the manufacture’s instruction (*supper*Golgi Kit, Bioenno Tech, LLC, Cat# 003010). After impregnation, sections (100 to 200 µm) were obtained using a vibratome and the sections were mounted on gelatin-coated glass slides, and stained ^135^. Images were taken by using a Zeiss Imager II deconvolution microscope with SlideBook 6.0 software. For quantification of spines, images were acquired as a series of z-stack at 0.1-µm step to create sequential images enabling spine counting, and spine morphology measurements on 3D images using a 100 X oil objective. NeuronStudio (Version 0.9.92; http://research.mssm.edu/cnic/tools-ns.html, CNIC, Mount Sinai School of Medicine) was used to reconstruct and analyze dendritic spines as described previously ^37,53,70^.

### Transmission electron microscopy (TEM)

For TEM experiments, all tissue were processed using freshly prepared solutions on the day of perfusion, as described previously ^136^. Briefly, animals were anesthetized and transcardially perfused with normal saline followed by 2.5% glutaraldehyde/4% paraformaldehyde EM fixative (dissolved in 0.16 M NaH_2_PO_4_/0.11M NaOH buffer, pH 7.2-7.4) for 30 min. After perfusion, entire mouse carcasses were post-fixed for at least 1 week in the same EM fixative. The hippocampus was then dissected out and incubated overnight in 0.1M sodium cacodylate buffer, followed by incubation in a 2% OsO_4_ solution and gradient ethanol dehydration. Samples were incubated in propylene oxide, left in 100% PolyBed resin for 36 hours, and embedded in flat molds at 55°C for 36 hours. After embedding, the molds were processed, sectioned at a thickness of 90 nm, and imaged on a JEOL 1400 electron microscope in the Electron Microscopy Lab at UT Health San Antonio. Synapses were identified by the presence of synaptic vesicles and postsynaptic densities. The number of synapses was manually accounted and quantified in each image.

### AAV injection

WT or *mgll*^loxp/loxp^ mice at 2 months of age were anesthetized with ketamine/Xylazine (200/10 mg/kg) and placed in a stereotaxic frame. AAV9-hsyn-eGFAP-cre.WPRE or AAV9-hsyn-eGFAP vectors were provided by Addgene (Watertown, MA). AAV Vectors were stereotaxically injected into both sides of the hippocampus in WT or *mgll*^loxp/loxp^ mice at the coordinate: AP, -2.3, ML, 2, and DV, -2, as described previously ^34,40,54,70^. The behavioral performance, including novel object recognition and the Morris water maze tests, was carried out 30 days after injection of vectors.

### BrdU labeling and immunofluorescence

Immunofluorescence analysis was performed to assess neurogenesis in coronal brain sections from WT and mgll KO mice. BrdU (10 mg/kg) was intraperitoneally injected into the mice every 2 hours, three times a day for two days. Six days after the injections, the animals were anesthetized with ketamine/xylazine (200/10 mg/kg) and subsequently transcardially perfused with PBS, followed by 4% paraformaldehyde in phosphate buffer. The brains were quickly removed from the skulls, fixed in 4% paraformaldehyde overnight, and then transferred into PBS containing 30% sucrose until they sank to the bottom of the small glass jars. Cryostat sectioning was performed on a freezing vibratome at 40 μm, and a series of five equally spaced sections (every 10 sections) were collected in 0.1M phosphate buffer. Free-floating sections were immunostained using BrdU and doublecortin (DCX) antibodies, followed by incubation with the corresponding fluorescent-labeled secondary antibody. 4′,6-Diamidino-2-phenylindole (DAPI), a fluorescent stain that binds strongly to DNA, was used to detect cell nuclei in the sections. Immunofluorescence imaging was conducted using a Zeiss deconvolution microscope with Slidebook software 6.0 (Intelligent Imaging Innovations, Denver, Colorado), as described previously ^34,40^.

### Single-cell/nucleus sample preparation

Single-cell suspensions from WT, mgll tKO, aKO, and nKO mice were prepared using an Adult Brain Dissociation Kit (MACS Miltenyi Biotec, Cat# 130-107-677) according to the manufacturer’s instructions, with some modifications, as previously described ^16,34^. Single-nucleus suspensions from WT, tKO, aKO, and nKO mice were prepared using a protocol previously described ^137^ with some modification. Briefly, frozen hippocampi from five animals for each genotype were homogenized in ice-cold lysis buffer on ice. The resulting suspension was filtered through a 20 μm filter to remove debris and centrifuged at 500 g for 5 minutes at 4°C. Nuclei were washed and filtered twice with nuclei wash buffer. Finally, the pellets were carefully resuspended in a suitable volume of nuclei buffer to achieve a concentration of 500-1,000 nuclei/μl for subsequent capture, as previously described ^137^.

### Single-cell/nucleus RNA sequencing library preparation

Single-cell or nucleus suspensions were loaded into the 10x Genomics Chromium microfluidic chips with the intention of capturing 8,000 to 10,000 cells within individual Gel Beads-in-emulsion (GEM). Within the GEMs, cell lysis occurred, and RNA was reverse transcribed using poly(dT) priming, during which Cell Barcodes and Unique Molecular Identifiers (UMIs) were incorporated into the cDNA. The prepared libraries, following the 10x Genomics 3’ Gene Expression v3 protocol, were sequenced using the Illumina NovaSeq 6000 system at the Genome Sequencing Facility (GSF) of Greehey Children’s Cancer Research Institute at UT Health San Antonio.

### Single-cell/nucleus RNA-seq data analysis

ScRNA-seq data were analyzed using the Cell Ranger software suite (v4.0) and the Seurat R package (v5.0.3) in R (v4.2.1), as described previously ^16,34,85,138^. Cells or nuclei with mitochondrial gene expression over 30% or ribosomal gene expression over 20% were deemed low quality and excluded. Genes appearing in fewer than 10 nuclei were discarded. Filtering parameters set limits of 400 to 6000 genes per nucleus and a maximum of 35,000 UMIs. Normalization of feature expression was performed using the NormalizeData function with a scale factor of 10,000. The most variable features were identified using the FindVariableFeatures function, selecting 5000 features via the ’vst’ method. Data scaling was performed using ScaleData. Dimensionality reduction was conducted using PCA on these 5000 variable genes via the RunPCA function, and the top 25 principal components were used for clustering nuclei with a resolution of 2. Visualization was achieved with uniform manifold approximation and projection (UMAP).

Data from the snRNA-seq database and the scRNA-seq database were integrated using the Seurat package. Comparisons were made between snRNA-seq data with other snRNA-seq data, and scRNA-seq data with other scRNA-seq data. If snRNA-seq data were found in non-neuronal clusters or scRNA-seq data in neuronal clusters, they were removed. Batch effects were not removed at this step since snRNA-seq data were not compared with scRNA-seq data.

For cell type classification, at least two specific markers were used to confirm cell identity. Syt1 and Rbfox3 were used as neuronal markers. Excitatory neurons were identified based on the expression of Nrgn and Slc17a7, while Gad1 and Gad2 were used as specific markers for inhibitory neurons. For glial cells, Aqp4 and Gja1 were used to identify astrocytes, while Aif1 and Tmem119 were used to identify microglia. Additionally, Plp1 and Mog were used for oligodendrocyte lineage cells, Cldn5 and Cdh5 for endothelial cells, Pdgfrb and Acta2 for pericytes, and Dcx and Prox1 for immature neurons.

Differentially expressed genes (DEGs) in each genotype were identified using the FindMarkers function of the Seurat package in R. This process computed average log2 fold changes, the percentage of cells expressing each gene in the groups compared, and provided P values and adjusted P values calculated using the Wilcoxon method. Synaptic genes were downloaded from the SYNGO (Synaptic Gene Ontologies) website. Genes related to adult neurogenesis were downloaded from the MANGO (Mammalian Adult Neurogenesis Gene Ontology) website. The ‘inner_join’ function in the ‘dplyr’ R package was then used to filter DEGs based on the downloaded genes. DEGs were considered significant if they had P < 0.05 and a log2(fold change) greater than 0.1 or less than -0.1.

Heatmaps, volcano plots, violin plots, and GO analyses were generated using the clusterProfiler R package and OriginLab 2024. Violin plots were created using the dittoSeq (1.8.1) package in R (4.2.1) ^139^.

### Novel object recognition test

The novel object recognition (NOR) test was performed to assess memory retention as described previously ^54,70^. Briefly, animals were first allowed to acclimate to the testing environment (habituation). The test included two stages: training and testing. In the first stage of the test, the animal was confronted with two identical objects, placed in an open field, and in the second stage, the animal was exposed to two dissimilar objects placed in the same open field: one familiar object, used in the first phase, and the other novel object. Exploration of an object was defined as time spent with the head oriented towards and within two cm of the object. The time spent exploring each of the objects in stage two was detected using the EthoVision video tracking system (Noldus). The recognition index (RI) was calculated based on the following equation: RI =T_N_/T_N_+T_F_), where T_N_ is the exploration time devoted to the novel object and T_F_ is the exploration time for the familiar object, as described previously ^140^.

### Morris water Maze test

The classic Morris water maze (MWM) test was used to determine spatial learning and memory, as described previously ^34,37,39–41,53,54^. A circular water tank (diameter 120 cm and 75 cm in high) was filled with water and the water was made opaque with non-toxic white paint. A round platform (diameter 15 cm) was hidden 1 cm beneath the surface of the water at the center of a given quadrant of the water tank. WT and MAGL KO mice that were treated with sham or TBI received learning acquisition training in the Morris water maze for 7 days and each session consisted of 4 trials. For each trial, the mouse was released from the wall of the tank and allowed to search, find, and stand on the platform for 10 seconds within the 60-second trial period. For each training session, the starting quadrant and sequence of the four quadrants from where the mouse was released into the water tank were randomly chosen so that it was different among the separate sessions for each animal and was different for individual animals. The mouse movement in the water pool was recorded by a video-camera and the task performances, including swimming paths, speed, and time spent in each quadrant, were recorded using an EthoVision video tracking system (Noldus, version 14). A probe trial test was conducted 24 hours after the completion of the learning acquisition training. During the probe test, the platform was removed from the pool, and the task performances were recorded for 60 seconds.

### Quantification and statistical analysis

Data are presented as mean ± S.E.M. Unless stated otherwise, Student’s t test, one or two-way analysis of variance (ANOVA) followed by post-hoc tests were used for statistical comparison when appropriate. Differences were considered significant when P< 0.05.

## Materials availability

Mouse lines generated in this study are available upon reasonable request from the corresponding author.

## Data availability

Sing-cell/nucleus RNA sequencing data have been deposited at the NCBI Gene Expression Omnibus (GEO) under accession number “GSE178226” and “GSE274338” and are publicly available as of the date of publication. All other data reported in this paper will be shared by the lead contact upon request.

## Acknowledgement

This work was supported by National Institutes of Health grants RF1NS076815, R01MH113535, and RF1AG081362 (to C.C.) and by startup funds from UT Health San Antonio, Joe R. & Teresa Lozano Long School of Medicine (to C.C.). RNA sequencing data was generated in the Genome Sequencing Facility at UT Health San Antonio, which is supported by UT Health San Antonio, NIH Shared Instrument grant S10OD030311, and CPRIT Core Facility Award (RP220662). The authors also thank Ms. Anastassia R. Nelson for animal care.

## Author Contributions

C.C. conceived the project and designed the experiments; D.Z., J.Z., M.H., F.G., J.H. J.L., X.M., J.W., Y.C., and S.Q., and C.C. performed the experiments and/or analyzed the data; C.C. supervised the work and C.C. wrote the manuscript.

