## Supplemental figures for "Overabundant endocannabinoids in neurons are detrimental to cognitive function"

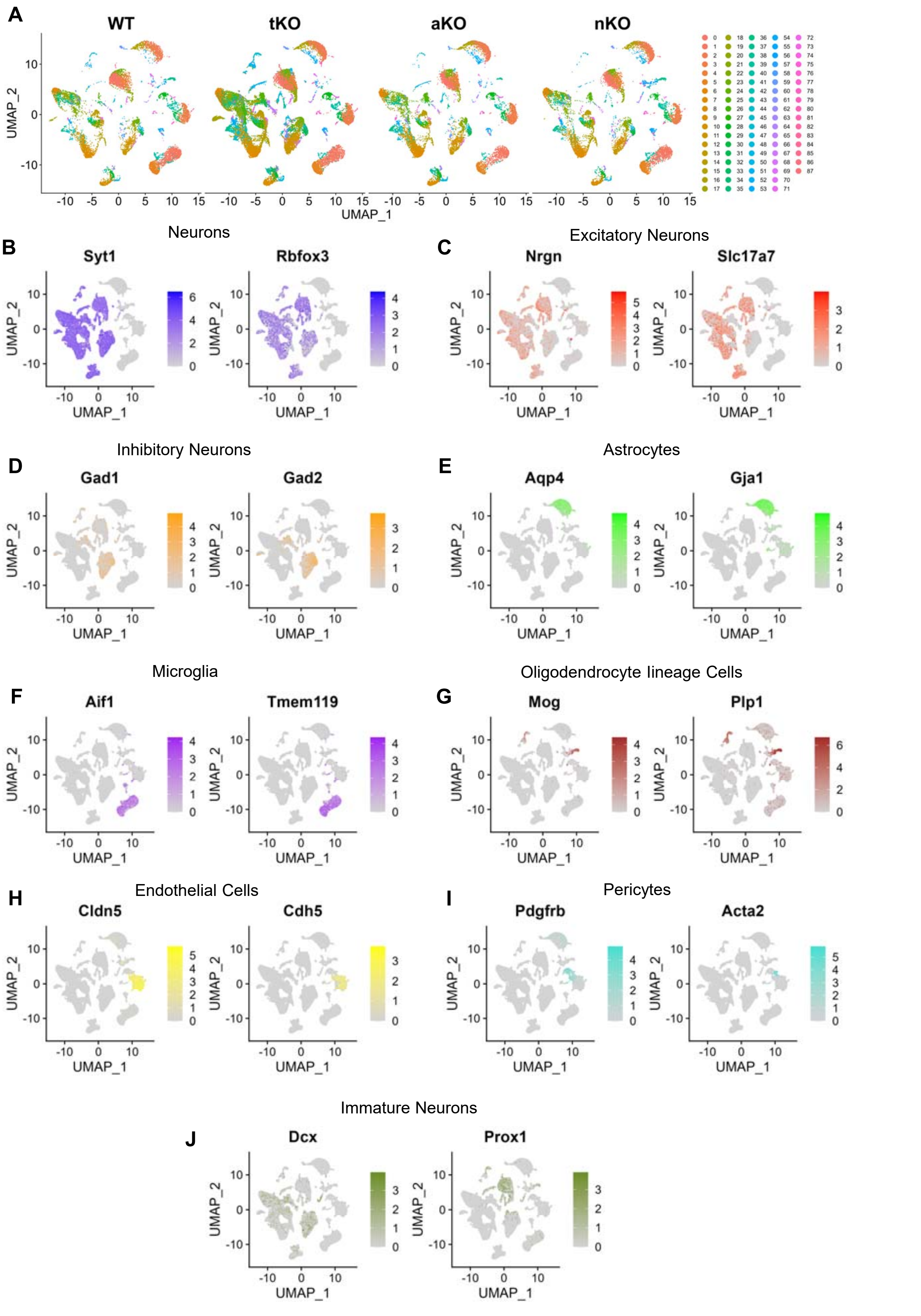


Figure-S1. Cell cluster annotation of single-nucleus and single-cell RNA-seq data from WT, tKO, aKO, and nKO mice. **A**. UMAP plots of snRNA-seq data from mouse hippocampi and scRNA seq-data from mouse hippocampi and cortex. Cells are colored by cluster assignments. **B-J**. Feature plot displaying the expression of marker genes used to identify main cell subsets: sy1/rbfox3 (neurons), nrgn/slc17a7 (excitatory neurons), gad1/gad2 (inhibitory neurons), aqp4/gja1 (astrocytes), aif1/tmem119 (microglia), plp1/mog (oligodendrocytes lineage cells), cldn5/cdh5 (endothelial cells), pdgfrb/acta2 (pericytes) and dcx/prox1 (immature neurons).

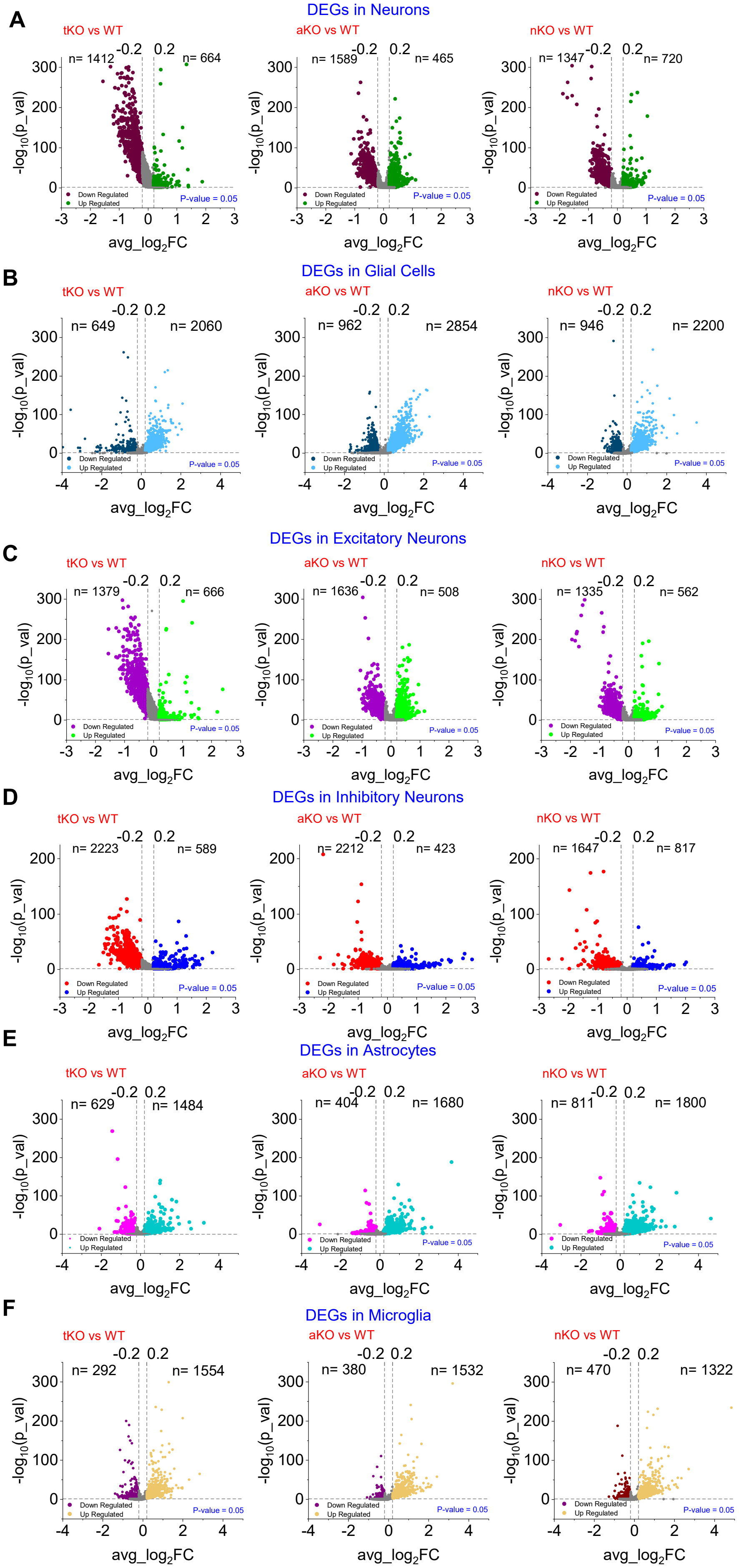


Figure-S2. Volcano plots representing the differential expression analysis of genes in neurons (**A**), glial cells (**B**), excitatory neurons (**C**), inhibitory neurons (**D**), astrocytes (**E**) and microglia (**F**) across tKO, aKO, and nKO mice.

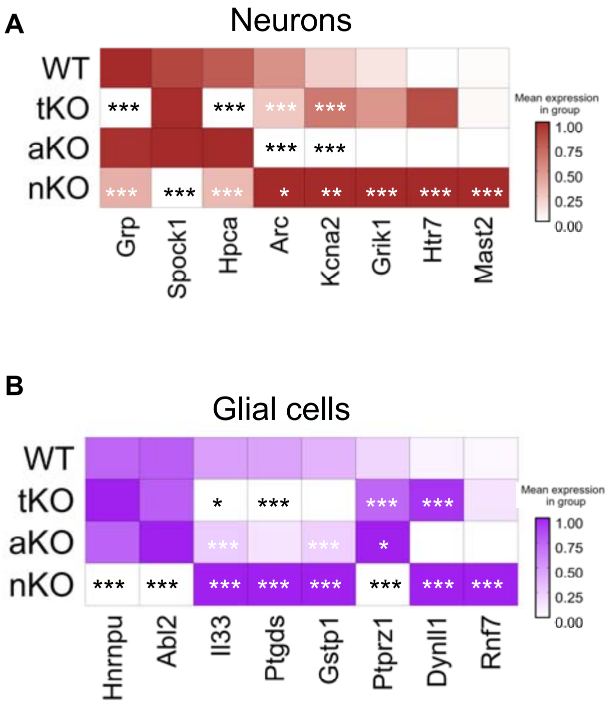


Figure-S3. Heatmaps for representative genes associated with synaptic function in neurons (**A**) and glial cells (**B**). *P<0.05; **P<0.01; ***P<0.001 compared with WT.
